# Impact of different synonymous codon substitution strategies on SARS-CoV-2 nucleocapsid protein expression in *Escherichia coli*

**DOI:** 10.1101/2024.11.06.622014

**Authors:** Adebayo J. Bello, Ayomide O. Omotuyi, Olajumoke B. Oladapo, Adegbayi A. Adekunle, Kingsley U. Udechime, Olufunke T. Olugbenga, Abdulzamad B. Akinwande, Adeyemi F. Odewale, Eban Davis, Olawale Adamson, Onikepe Folarin, Joy Okpuzor, Joseph B. Minari

## Abstract

Synonymous codon substitution, a gene engineering approach in synthetic biology, has been effective in improving the codon composition of recombinant genes of interest based on various criteria without altering the amino acid sequence. The SARS-CoV-2 virus nucleocapsid (N) protein is a stable, conserved and highly immunogenic that is less prone to mutation during infection, making it a key antigen in *in vitro* diagnosis, vaccine development, immunological and structural studies. While reports have focused on applying optimized N protein for different applications, the basic parameters used by different optimization tools for choosing the best approach for the N gene synonymous codon substitution are often neglected. Here, we analyzed the influence of different synonymous codon substitution strategies on SARS-CoV-2 N-protein expression in *E. coli*. Using different codon optimization (CO) and harmonization (CH) tools, we predicted and compared how parameters such as GC content, Codon Adaptation Index, codon quality and number of rare codons present in these sequences affect the N-protein expression. Our results also show that Minimum Free Energy (MFE) and RNA structure of N-term and C-tail of the N-protein coding sequence influence protein folding. We then predicted that the SR-rich region of the N-protein may contribute to slowing down the elongation rate during translation. This work presents a fundamental analysis of how different optimization tools affect SARS-CoV-2 N-protein expression and folding and suggests a basic approach to choosing the best strategy for optimal expression and folding of the protein for further studies.

**Graphical Abstract:** 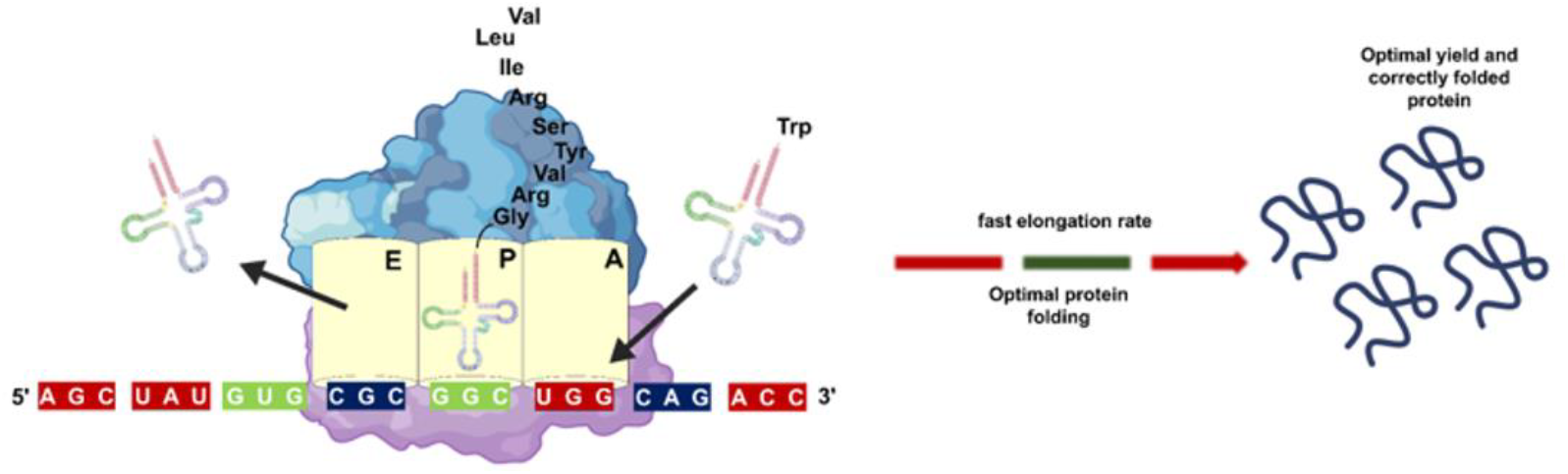

**Highlights:** Codon substitution affects SARS-CoV-2 nucleocapsid (N) protein expression.

SARS-CoV-2 N-protein expression varies with different codon optimization tools.

RNA structures of N- and C-term impact RNA stability, protein expression and folding.

SR-rich region of the N-protein may slow down elongation rate during translation.

## Introduction

Severe acute respiratory syndrome coronavirus (SARS-CoV-2) is a fast-transmissible virus among humans, and it is responsible for coronavirus disease (COVID-19) [1]. Since the outbreak of the disease in the city of Wuhan, Hubei Province, China in December 2019, there have been more than 700 million confirmed cases and over 7 million deaths globally as of July 2025. (https://www.worldometers.info/coronavirus/). This disease poses an incomparable threat to global public health, and new theranostic discoveries are needed to combat the disease. Despite new vaccine development and drug candidates that provide a saving route to reduce the burden of the coronavirus disease, there is a need for continuous studies on the virus and its effect on host-pathogen interaction.

The SARS-CoV-2 nucleocapsid (N) protein is one of the four structural proteins with 419 bp amino residues. The N-protein is the most abundant, relatively conservative protein in coronaviruses [2], and it is one of the most studied proteins because of its importance as an antiviral target [3]. While serving as RNA binding proteins (RBPs), the N-protein is responsible for packaging the viral RNA genome into ribonucleotide complex (RNP) through protein-RNA interaction [4], determines pathogenicity and virulence [5], regulates progression of the host cell cycle, plays an important role in interaction of the virus with the host, as well as the apoptosis [6]. The protein has been reported to be highly expressed during infection [7]. These characteristics make the protein a key factor in the viral life cycle.

Most studies on SARS-CoV-2 involve expression of the viral proteins either *in vitro* or *in vivo* to develop drug candidates, vaccines, and diagnostic biomolecules to provide safe working environments [8], [9] [10]. Likewise, *in silico* studies have formed an integral part of SARS-CoV-2 research either by designing new bioinformatics tools or making use of existing algorithms to understand the virus structural architecture, design new drugs and vaccines against the virus [11], [12], [13]. However, for efficient viral protein expression, it has been proven that synonymous codon substitution [14] is an important factor to consider when expressing the protein in a heterologous host such as *Escherichia coli*. Because multiple codons can code for the same amino acid because of the genetic code degeneracy, which can thus influence protein expression, preferential usage of codons termed codon usage bias however varies in organisms [15]. Thus, different organisms are codon-biased during protein expression based on the abundance of corresponding intracellular aminoacyl-tRNAs during translation [16].

Production of synthetic nucleic acids and recombinant proteins is common in most laboratories for research purposes and industrial applications. One strategy that has improved this process over time is optimizing codons of genes of interest to fit into the codon usage bias of the heterologous host [17], [18]. This process of codon optimization or synonymous codon substitution for efficient production of recombinant proteins in heterologous hosts is one of the major bases of modern biotechnology. This approach involves engineering gene sequences to enhance protein expression without changing the amino acid letters and employs translation elongation rate as a means of improving the expressed protein.

Different tools have been designed to improve protein expression in heterologous hosts. Some reports have also documented the codon usage pattern and codon bias of the SARS-CoV-2 genome [19], [20], [21] while the virus N-protein optimization and expression for different downstream studies such as diagnostic [22], vaccines [23], immunological assessment [24], and structural studies [25] have also been reported. However, while these reports focused on applying optimized protein, the basic parameters for synonymous codon substitution used by different optimization tools are often neglected. The unreported comparison which usually leads to differences in the protein yield, solubility, and folding may be critical to SARS-CoV-2 nucleocapsid proteins for vaccine production, diagnostics, and structural and immunological studies. Thus, determining the most suitable tool for desirable optimized protein is vital, as different tools have unique parameters for optimizing nucleotide sequences [26]. Also, no report to the best of our knowledge has investigated how SAR-CoV-2 N-protein expression is affected by codon harmonization, a strategy to adapt the gene of interest into heterologous for optimal protein folding [27].

In this study, we performed synonymous codon substitution and compared the underlining parameters affecting SARS-CoV-2 nucleocapsid protein expression using JCat and Genewiz codon optimization algorithms, and codon harmonization tool. These optimization tools were chosen based on frequent usage for protein expressions in reported studies [27], [28], [29]. Overall, the genetic code substitution strategy predicted variation in expression analysis which may affect the protein yield and folding properties.

## 2.1. Methods

### 2.1. Retrieval of Gene Coding Sequences from Database

The gene coding sequence for SARS-CoV-2 nucleocapsid (N) protein was obtained from nucleotide database of National Center for Biotechnology Information (NCBI) website www.ncbi.nlm.nih.gov/nuccore/NC_045512.2?from=28274&to=29533&report=fasta). The FASTA format of nucleotide sequence from 28274-29533 bp with Gene ID; 43740575 was used for this study.

### 2.2. Synonymous codon substitution strategies

Two codon optimization algorithms (JCat and Genewiz) and one codon harmonization algorithm [29] were used for the synonymous codon substitution of the nucleocapsid gene sequence. The coding sequence for the N protein (wild type) was obtained from the NCBI database and pasted into each algorithm while selecting *Escherichia coli* as the heterologous expression host for the N protein. Then, the tools were used to generate different variants of the N protein coding sequence (Suppl. File 1). All other parameters were set at default.

### 2.3. *In silico* analysis of the nucleocapsid gene codon sequence

Analysis of the gene coding sequences of the wildtype and the variants for the GC content, Codon Adaptation Index (CAI), rare codons frequency, and codon quality class were calculated on the Biologics Corp’s Bioinformatics Center (https://www.biologicscorp.com/tools/) using *E. coli* as a heterologous host. The rare codon clustering of the substituted synonymous codons was calculated using the percentage MinMax Calculator [30].

### 2.4. Cloning and protein expression

All the *E. coli* cells were grown and maintained in Luria-Bertani (LB) medium with constant shaking at 220 rpm at 37^°^C. Routinely, 25 µg/ml chloramphenicol (Cm) and 50 μg/ml kanamycin (Km) when added when necessary. The different optimized sequences were flanked by NdeI and XhoI restriction enzymes and synthesized by Integrated DNA Technologies (IDT). The cDNA sequence was cloned into PET28a+ derivative plasmid between NdeI and XhoI sites. NEB10 *E. coli* strain was transformed with the plasmids pJB30_JCat, pJB31_Genewiz, and pJB33_CH through electroporation. The clones were selected on plates containing both Cm and Km and grown in liquid culture overnight at 37^°^C. Plasmid DNA was isolated from the overnight culture using QIAprep^®^ Spin Miniprep Kit (Qiagen) and the precision and orientation of the sequence were confirmed by sequencing.

BL21(DE3)pLysS *E. coli* strains were chemically transformed with the plasmids and grown overnight in media containing Cm and Km. Expression of the transformed cells with relevant plasmids was carried out under the control of an inducible T7 promoter, as described in [31]. Fresh overnight culture was added to a sterile LB medium, and the cells were induced with IPTG at a final concentration of 1 mM after 100 minutes of initial growth and further incubated for 135 minutes post-induction before harvesting. Cell pellets from 5ml culture were first suspended in 10mM Tris-HCl pH 7.5 before adding 500 µl CelLytic® B Lysis buffer (Sigma, UK). The mixture was vortexed briefly and incubated at room temperature with constant shaking (60 rpm; 5 minutes) before centrifuging (13,000 rpm; 5 minutes). Then, protein loading samples from the supernatant were prepared in 4X Laemmli buffer containing 50mM final concentration of Dithiothreitol (DTT). The sample mixture was incubated at 95^°^C for 5 minutes and spun down shortly. Sodium dodecyl-polyacrylamide gel electrophoresis (SDS-PAGE) was used to separate the protein bands on a 4–20% Mini-PROTEAN® TGX Stain-Free™ Protein Gels (Bio-Rad). The gel image was visualized and analyzed using ChemiDoc™ Imaging System (Bio-Rad) and ImageJ software v1.5j respectively.

### 2.5 RNA stability

The RNA stability of the N terminal region of the sequences was determined using RNAfold [32]. The secondary structures of their RNA were generated in FORNA format, and the Minimum free energy (MFE) was generated for each sequence.

### 2.6. Statistical analysis

All descriptive values were calculated using Microsoft Excel version 2402 (2024). Significant differences were calculated using one-way ANOVA where ^*^P<0.05.

## 3. Results and discussion

As one of the basic principles in synthetic biology, codon optimization can conveniently enable high expression of toxic proteins in microbial hosts [33]. As the research on SARS-CoV-2 continues to evolve, it is important to determine the best-fit parameters for expressing its toxic protein in microbial hosts.

### 3.1. Prediction of the effect of different synonymous codon substitution strategies

In this study, we compared the impact of different codon optimization strategies on the expression of SARS-CoV-2 nucleocapsid protein in *E. coli*. Using *In Silico* analysis, we showed that codon optimization is an important tool for SARS-CoV-2 N-protein and compared the different parameters responsible for increased protein expression. The optimized sequences differ from the wildtype by 24.44, 26.98, and 23.97 % for JCat, Genewiz, and CH optimized sequence respectively as calculated from the Percentage Index Matrix (Supplemental Figure S1) after performing sequence alignment using ClustalW. The percentage differences between the optimized sequence and the wildtype sequence show that the codon substitution strategy has significant impact on the protein expression.

The presence of rare codons can lead to a reduction in the protein yield in microbial hosts. Several reports have shown that too many rare codons do not only delay translation, but they can contribute to early termination leading to production of premature proteins [34], [35], [36], [37]. Our result shows that the low rare codons in the Genewiz optimized sequence can contribute to increased elongation process during translation, as against the low frequency JCat, and CH (Fig 1a). Most of the optimized codons also fall within the codon quality class of 91-100 (Fig 1b), indicating a strong influence on the N-protein expression in *E. coli*.

**Fig 1.**
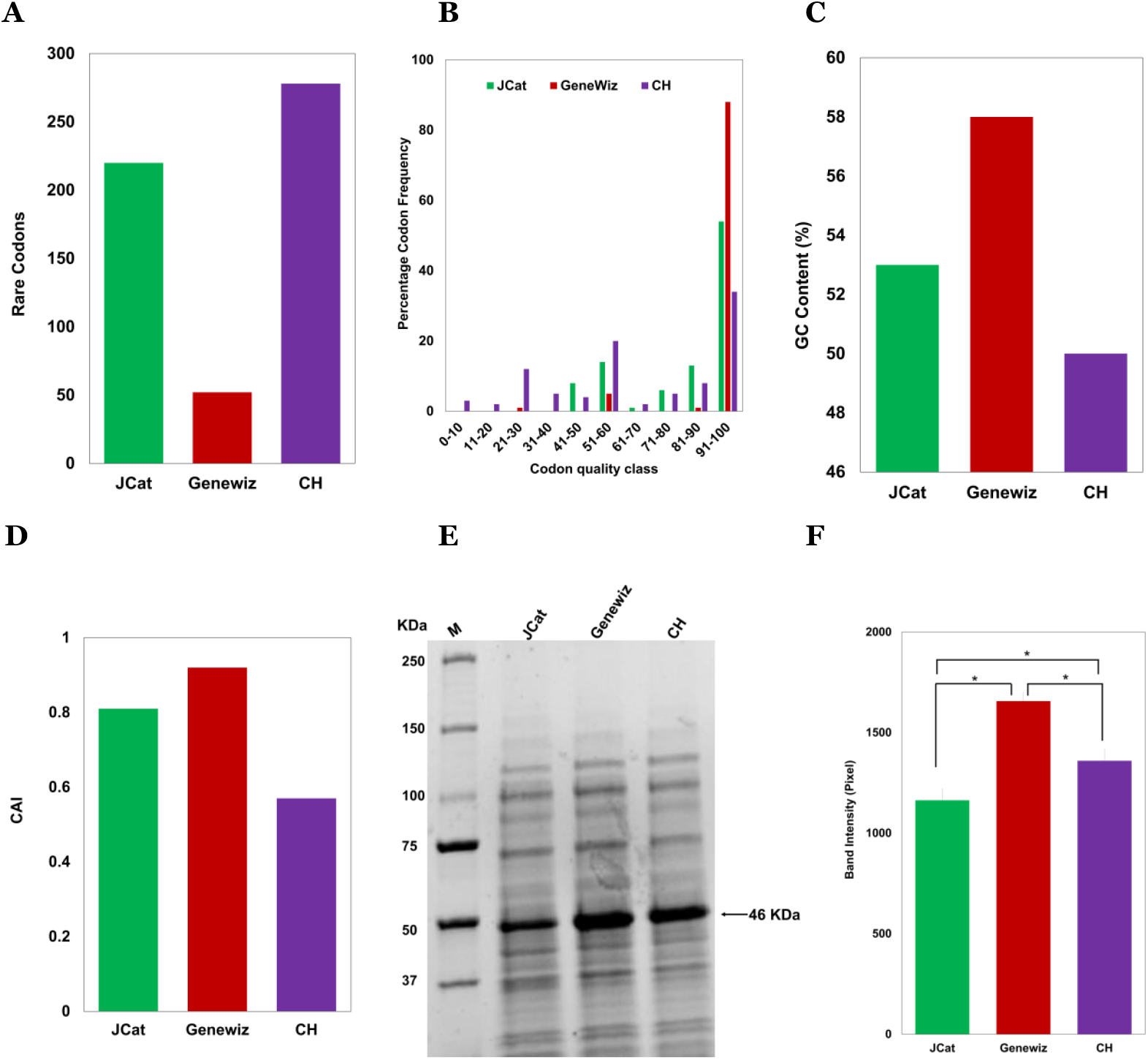
Effects of different parameters influenced by the genetic code substitution. (A) Rare codon. (B) Codon quality class. (C) percentage GC content. (D) Codon Adaptation Index. (E) gel image of the expressed protein, M – protein marker. (F) average band intensity and standard error bar calculated from the gel image in triplicate. The asterisk shows significant at P<0.05.

The calculated GC contents for the optimized sequences are 53, 58, and 50% for JCat, Genewiz, Geneart, and CH respectively (Fig 1c). The CAI are 0.81, 0.92, and 0.57 for JCat, Genewiz, and CH respectively (Fig 1d). GC content and CAI have been good predictive parameters for increased protein expression. Reports have shown that higher GC content can increase mRNA levels [38] and influence protein expression [39], [40]. Although some reports argued that CAI is not a comprehensive measure to determine the level of protein expression, however, Lin *et al*., (2022) showed that an increase in CAI is a good predictive value for experimental approach [41]. In our study, Genewiz optimized sequence showed the highest values for the calculated parameters. Generally, the optimized sequence influenced the predictive parameters; increased the GC content and CAI, reduced rare codons frequency, and improved codon quality class compared to JCat and CH. Fulfilling all these parameters positively, we suggest the possibility of higher expression of the SARS-CoV-2 N-protein in *E. coli* using Genewiz codon optimization tool.

### 3.2. protein expression

To validate the experiment, we cloned and expressed the plasmids carrying each coding sequence in BL21(DE3)pLysS *E. coli* cells. The cloned codon-optimized genes flanked by NdeI and Xhol were confirmed after sequencing to show the correct sequence and their orientation. The N-protein expressed after IPTG induction (Fig 1E) shows that the synonymous substitution to the coding sequence did not affect the cell growth condition by inducing toxicity [42], [43]. The Genewiz recombinant N-protein was more highly expressed than JCat and CH (Fig 1F) using protein band quantification. However, the expression of JCat codon optimization was significantly lower than CH by forming inclusion bodies [44] despite fulfilling all the *in-silico* analyses positively and having the highest number of relative codon frequency (Table 1).

**Table 1:** codon usage frequency for each optimized sequence calculated from https://www.biologicscorp.com/tools/CodonUsageCalculator

|  |  | Jcat |  | Genewiz |  | CH |  |
| --- | --- | --- | --- | --- | --- | --- | --- |
| Amino acid |  | Codon | Frequency | Codon | Frequency | Codon | Frequency |
| Ala | A | GCU | 1 | GCG | 0.89 | GCC | 0.38 |
| Arg | R | CGU | 0.93 | CGC | 0.52 | AGA | 0.41 |
| Asn | N | AAU | 1 | AAC | 0.86 | AAC | 1 |
| Asp | D | GAC | 1 | GAU | 0.96 | GAC | 1 |
| Cys | C |  |  |  |  |  |  |
| Glu | E | GAA | 1 | GAA | 1 | GAG | 1 |
| Gln | Q | CAG | 0.97 | CAG | 0.54 | CAA | 0.8 |
| Gly | G | GGU | 1 | GGC | 0.74 | GGG | 0.93 |
| His | H | CAC | 1 | CAU | 1 | CAC | 0.75 |
| Ile | I | AUC | 1 | AUU | 1 | AUA | 1 |
| Leu | L | CUG | 0.96 | CUG | 0.93 | CUA | 0.85 |
| Lys | K | AAA | 0.77 | AAA | 0.84 | AAG | 1 |
| Met | M | AUG | 1 | AUG | 1 | AUG | 1 |
| Phe | F | UUC | 1 | UUU | 0.83 | UUC | 1 |
| Pro | P | CCG | 1 | CCG | 0.96 | CCC | 0.36 |
| Ser | S | UCU | 1 | AGC | 0.81 | UCU | 0.41 |
| Thr | T | ACC | 1 | ACC | 0.84 | ACA | 0.53 |
| Trp | W | UGG | 1 | UGG | 1 | UGG | 1 |
| Tyr | Y | UAC | 1 | UAU | 1 | UAC | 1 |
| Val | V | GUU | 1 | GUG | 0.75 | GUA | 0.5 |
| STOP |  | UAA |  | UAA |  | UAA |  |
| Average |  |  | 0.98 |  | 0.87 |  | 0.79 |

### 3.3. Impact of RNA structure

To further analyze the impact of synonymous codon substitution we predicted the secondary RNA structure to evaluate how they affect N-protein expression and folding. RNAs fold into 3D structures that range from simple helical elements to complex tertiary structures and quaternary ribonucleoprotein assemblies, and the functions of many regulatory RNAs depend on how their 3D structure changes in response to a diverse array of cellular conditions [45]. In this study, RNA secondary structure analysis of the N-term and C-tail (Fig 2A) was carried out to evaluate the structural differences and minimum free energy (MFE). The three RNA sequences have structures with different numbers of hairpin loops at the N-term (Fig 2B-D) and C-tail (Fig 2F-H). The “linear” structure of RNA in Genewiz at the N-term and C-tail may contribute to its stability and high protein expression, which agrees with Leppek et al., (2022) [46], that reducing the number of hairpin loops can enhance mRNA stability and increase protein expression. The linear pattern in Genewiz suggests an optimal translation rate to allow the N-term of the protein to fold properly before the C-terminal part is formed [47]. Conspicuously, the JCat optimized sequence resulted in an MFE of -41.6 kcal/mol, while Genewiz and CH optimized sequences yielded -40.2 and -5.4 kcal/mol respectively at N-term (Fig 1E). At the C-term, Jcat, Genewiz, and CH have -44.1, -38.0, and -23.7 kcal/mol respectively. The more negative the MFE, the more stable the RNA. However, highly stable RNA structure can interfere with ribosome access leading to low translation [48]. We interpreted this report to be the case of JCat with the highest MFE at both ends of the N-protein. Consequently, the Genewiz-optimized sequence appears to be the most advantageous choice in terms of RNA stability, considering the optimal MFE and the structural linearity.

**Fig 2.**
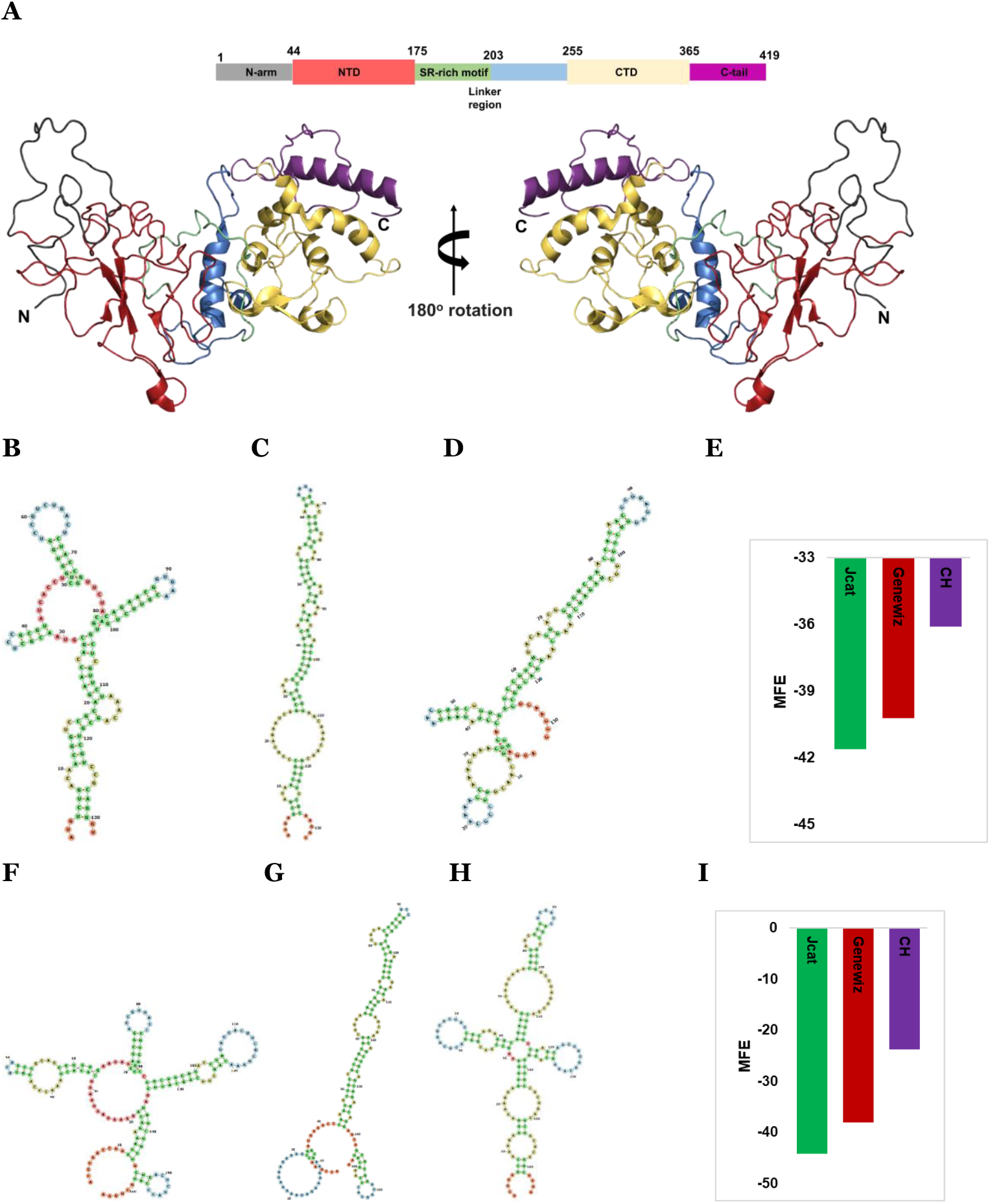
Genomic and structural arrangement of SARS-CoV-2 nucleocapsid protein and RNA. (A) top: schematic arrangement of nucleocapsid protein, highlighting the N-terminal domain (NTD) and C-terminal domain (CTD) flanking the SR-rich intrinsically disordered regions (IDRs). The NTD and CTD are flanked by other two IDRs, N-term and C-tail as illustrated. Down: The structure of the nucleocapsid protein (protein ID: 8FD5) visualized using PyMoL v3.0. RNA secondary structure of the N-term (132 nucleotide bases) for JCat (B), Genewiz (C), and CH (D), and minimum free energy (MFE) (E). RNA secondary structure of the C-term (165 nucleotide bases) for JCat (F), Genewiz (G) and CH (H), and minimum free energy (MFE) (I).

### 3.4. Percentage MiniMax analysis and protein folding

Similarly, throughout translation, the accurate pairing of codons and anticodons guides the incorporation of amino acids into the growing polypeptide chain. Adequate codon usage ensures that the translation machinery has access to a diverse pool of tRNAs, minimizing the occurrence of ribosomal pauses and misincorporations that could lead to protein misfolding [49]. Fig 3 shows the relative codon usage of the wildtype and optimized sequences using %MiniMax calculator with Genewiz showing a higher codon usage compared to the other sequences. The %MiniMax calculator, a freely accessible web-based portal and downloadable script, evaluates relative usage frequencies of synonymous codons within a protein sequence of interest [50]. They compare these results against a rigorous null model, providing valuable insights into codon usage patterns and optimization [30]. In Fig. 3A, the three sequences showed similar codon usage patterns, notably from amino residues 168 to 203 which may suggest similar folding patterns. The 36 amino acid residues majorly fall within the SR-rich IDR region of the nucleocapsid protein structure (Fig 3B). This may suggest the importance of serine and arginine bonds in protein expression and folding. However, the synonymous mutation resulted in 47.2% of the three RNA sequences (Fig 3C). Previous studies have experimentally confirmed the effect of codon substitution on protein folding [51], [52]. Liu et al 2021 [36] and Mauro and Chappell, 2014 [53] also provided extended reviews on the impact of codon optimization on protein folding. However, more experimental studies are needed to investigate the effect of SR-rich region and how synonymous mutation in the region may contribute to the overall protein expression and folding. Given these accounts, it is important to balance high protein yield with folding. Generally, the Genewiz pattern in this study may suggest higher protein expression and a well-folded protein.

**Fig 3.**
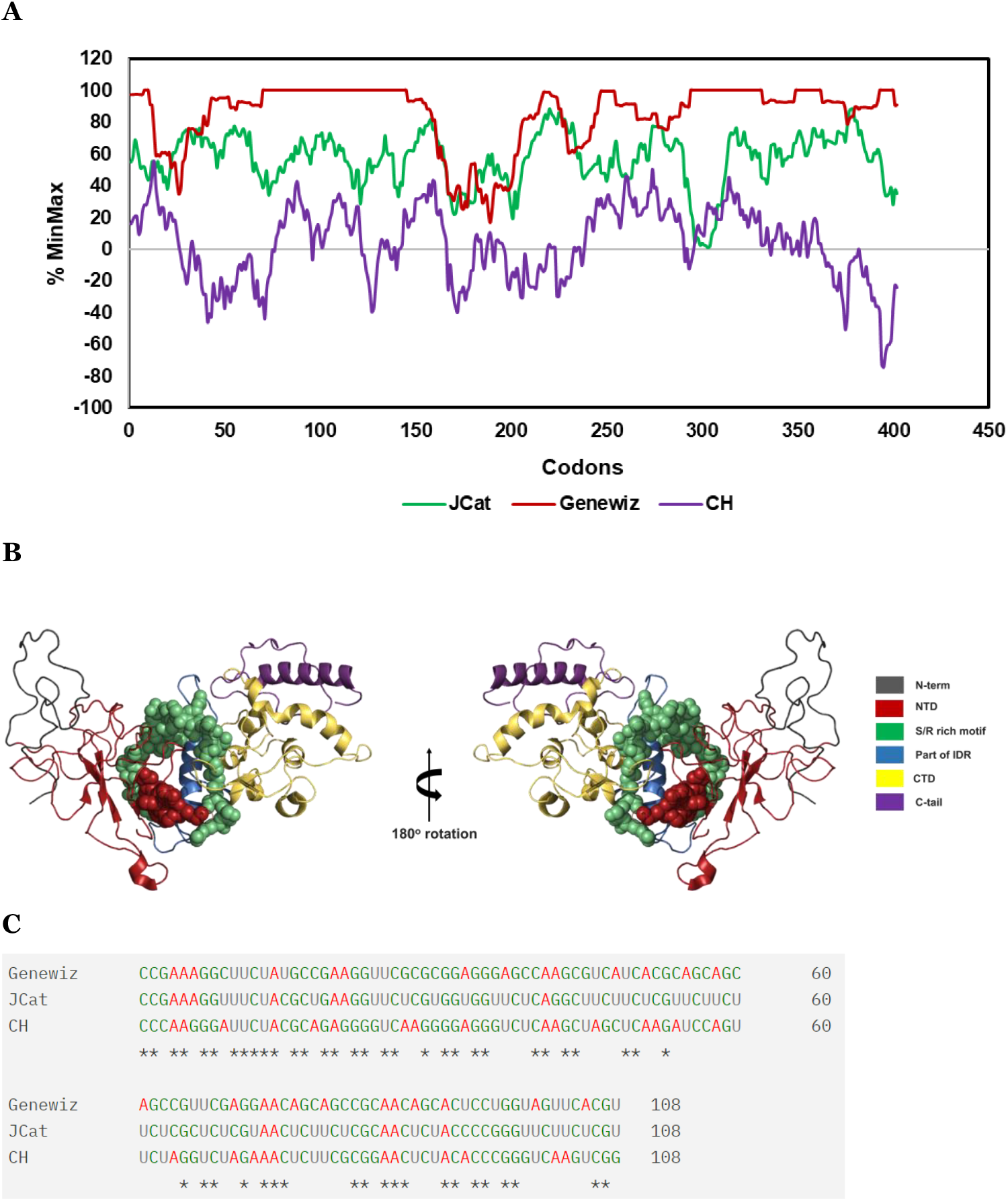
Predicted effect of the genetic code substitution on protein folding. (A) %MiniMax showing codon usage patterns of the three optimized sequences. (B) The structure of the nucleocapsid protein (protein ID: 8FD5) visualized using PyMoL v3.0. The spherical regions are SR rich region (green) and few residues P168 – E174 (red) upstream of the SR rich region. (C) sequence alignment of the optimized nucleotides at the regions in (B). The alignment was performed using ClustalW2 Multiple Sequence Alignment.

### 3.5. Elongation rhythm prediction

Synonymous code substitution finetunes codon bias usage to fit the abundance of tRNA available for protein production [54]. Therefore, we predict and represent how elongation rhythm is influenced by the three strategies used in this study (Fig 4). The decrease or increase in the translation rate influences the tRNA binding to the cognate mRNA as the ribosome performs its function which may eventually affect the protein yield and folding. A combination of “slow” and “rapid” tRNA:mRNA binding would generate a correctly folded protein but low yield (Fig4a). While a combination of “very rapid” and “rapid” with some “slow” tRNA:mRNA binding would generate optimal yield and correctly folded protein (Fig4b), an all through “very rapid” would lead to production of the high yield protein forming inclusion bodies (Fig4c)

**Fig 4.**
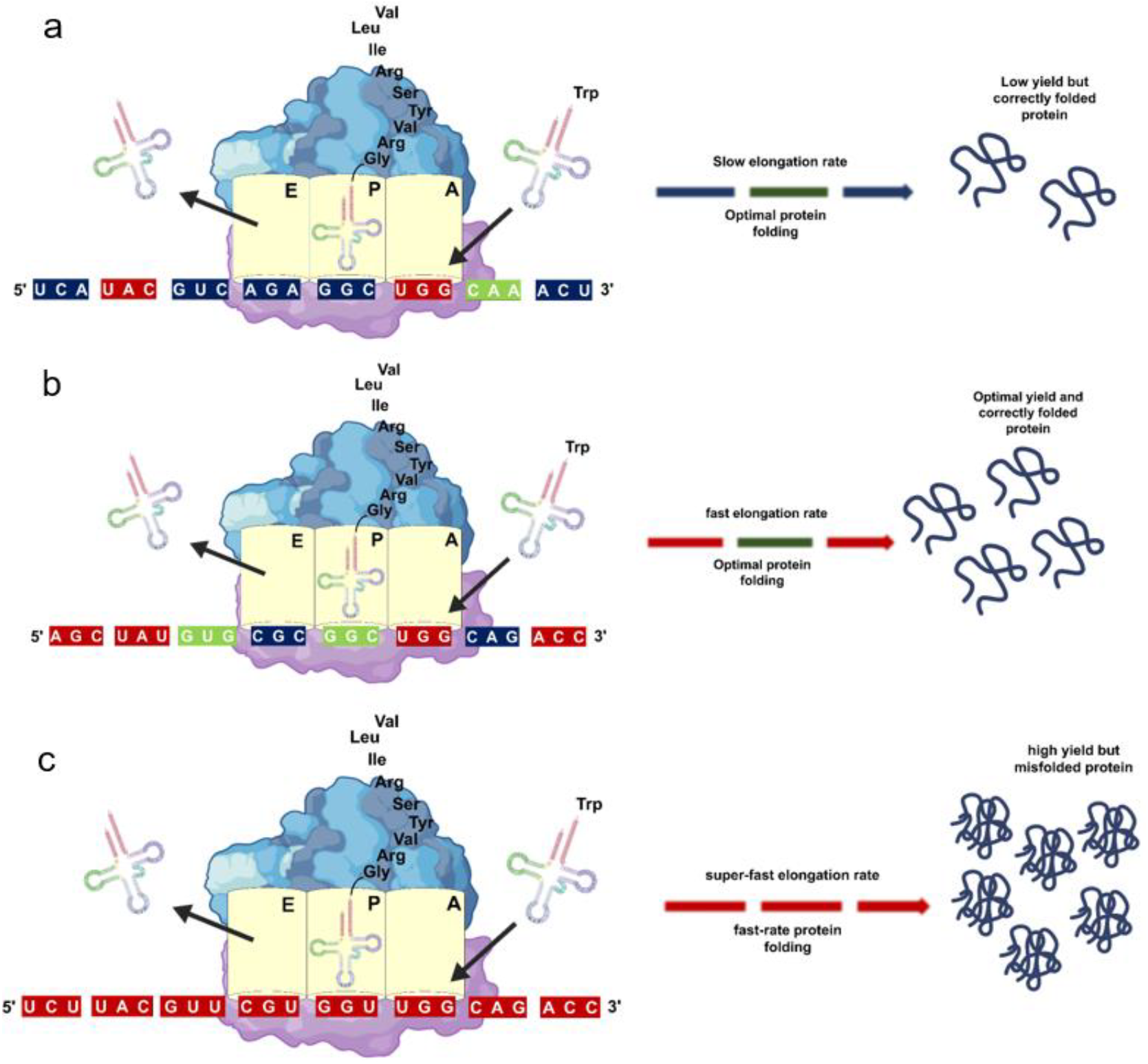
Effects of elongation rhythm pattern on the expression and folding of the protein from the three optimized tool. (A) codon harmonization (CH). (B) Genewiz. (C) JCat.

## Conclusion

This study shows that the optimized sequence of SARS-CoV-2 nucleocapsid protein for *E. coli* using Genewiz produced an optimal yield which could influence the folding of the protein compared to other optimized sequences. This work can provide a fundamental study for expressing viral proteins in microbial hosts to ascertain the best codon optimization strategies suitable for further studies such as designing new drugs and vaccine candidates, and structural studies.

