## Supplementary file for "Impact of different synonymous codon substitution strategies on SARS-CoV-2 nucleocapsid protein expression in *Escherichia coli*"

Nucleocapsid protein sequence

MSDNGPQNQRNAPRITFGGPSDSTGSNQNGERSGARSKQRRPQGLPNNTASWFTALTQHGKEDLKFPRGQ
GVPINTNSSPDDQIGYYRRATRRIRGGDGKMKDLSPRWYFYYLGTGPEAGLPYGANKDGIIWVATEGALN
TPKDHIGTRNPANNAAIVLQLPQGTTLPKGFYAEGSRGGSQASSRSSSRSRNSSRNSTPGSSRGTSPARM
AGNGGDAALALLLLDRLNQLESKMSGKGQQQQGQTVTKKSAAEASKKPRQKRTATKAYNVTQAFGRRGPE
QTQGNFGDQELIRQGTDYKHWPQIAQFAPSASAFFGMSRIGMEVTPSGTWLTYTGAIKLDDKDPNFKDQV
ILLNKHIDAYKTFPPTEPKKDKKKKADETQALPQRQKKQQTVTLLPAADLDDFSKQLQQSMSSADSTQA

N_WT_ref Seq

ATGTCTGATAATGGACCCCAAAATCAGCGAAATGCACCCCGCATTACGTTTGGTGGACCCTCAGATTCAA
CTGGCAGTAACCAGAATGGAGAACGCAGTGGGGCGCGATCAAAACAACGTCGGCCCCAAGGTTTACCCAA
TAATACTGCGTCTTGGTTCACCGCTCTCACTCAACATGGCAAGGAAGACCTTAAATTCCCTCGAGGACAA
GGCGTTCCAATTAACACCAATAGCAGTCCAGATGACCAAATTGGCTACTACCGAAGAGCTACCAGACGAA
TTCGTGGTGGTGACGGTAAAATGAAAGATCTCAGTCCAAGATGGTATTTCTACTACCTAGGAACTGGGCC
AGAAGCTGGACTTCCCTATGGTGCTAACAAAGACGGCATCATATGGGTTGCAACTGAGGGAGCCTTGAAT
ACACCAAAAGATCACATTGGCACCCGCAATCCTGCTAACAATGCTGCAATCGTGCTACAACTTCCTCAAG
GAACAACATTGCCAAAAGGCTTCTACGCAGAAGGGAGCAGAGGCGGCAGTCAAGCCTCTTCTCGTTCCTC
ATCACGTAGTCGCAACAGTTCAAGAAATTCAACTCCAGGCAGCAGTAGGGGAACTTCTCCTGCTAGAATG
GCTGGCAATGGCGGTGATGCTGCTCTTGCTTTGCTGCTGCTTGACAGATTGAACCAGCTTGAGAGCAAAA
TGTCTGGTAAAGGCCAACAACAACAAGGCCAAACTGTCACTAAGAAATCTGCTGCTGAGGCTTCTAAGAA
GCCTCGGCAAAAACGTACTGCCACTAAAGCATACAATGTAACACAAGCTTTCGGCAGACGTGGTCCAGAA
CAAACCCAAGGAAATTTTGGGGACCAGGAACTAATCAGACAAGGAACTGATTACAAACATTGGCCGCAAA
TTGCACAATTTGCCCCCAGCGCTTCAGCGTTCTTCGGAATGTCGCGCATTGGCATGGAAGTCACACCTTC
GGGAACGTGGTTGACCTACACAGGTGCCATCAAATTGGATGACAAAGATCCAAATTTCAAAGATCAAGTC
ATTTTGCTGAATAAGCATATTGACGCATACAAAACATTCCCACCAACAGAGCCTAAAAAGGACAAAAAGA
AGAAGGCTGATGAAACTCAAGCCTTACCGCAGAGACAGAAGAAACAGCAAACTGTGACTCTTCTTCCTGC
TGCAGATTTGGATGATTTCTCCAAACAATTGCAACAATCCATGAGCAGTGCTGACTCAACTCAGGCCTAA

N_JCat

ATGTCTGACAACGGTCCGCAGAACCAGCGTAACGCTCCGCGTATCACCTTCGGTGGTCCGTCTGACTCTA
CCGGTTCTAACCAGAACGGTGAACGTTCTGGTGCTCGTTCTAAACAGCGTCGTCCGCAGGGTCTGCCGAA
CAACACCGCTTCTTGGTTCACCGCTCTGACCCAGCACGGTAAAGAAGACCTGAAATTCCCGCGTGGTCAG
GGTGTTCCGATCAACACCAACTCTTCTCCGGACGACCAGATCGGTTACTACCGTCGTGCTACCCGTCGTA
TCCGTGGTGGTGACGGTAAAATGAAAGACCTGTCTCCGCGTTGGTACTTCTACTACCTGGGTACCGGTCC
GGAAGCTGGTCTGCCGTACGGTGCTAACAAAGACGGTATCATCTGGGTTGCTACCGAAGGTGCTCTGAAC
ACCCCGAAAGACCACATCGGTACCCGTAACCCGGCTAACAACGCTGCTATCGTTCTGCAGCTGCCGCAGG
GTACCACCCTGCCGAAAGGTTTCTACGCTGAAGGTTCTCGTGGTGGTTCTCAGGCTTCTTCTCGTTCTTC
TTCTCGCTCTCGTAACTCTTCTCGCAACTCTACCCCGGGTTCTTCTCGTGGTACCTCTCCGGCTCGTATG
GCTGGTAACGGTGGTGACGCTGCTCTGGCTCTGCTGCTACTGGACCGTCTGAACCAGCTGGAATCTAAAA
TGTCTGGTAAAGGTCAACAGCAGCAGGGTCAGACCGTTACCAAAAAGTCTGCTGCTGAAGCTTCTAAAAA
GCCGCGTCAGAAACGTACCGCTACCAAAGCTTACAACGTTACCCAGGCTTTCGGTCGTCGTGGTCCGGAA
CAGACCCAGGGTAACTTCGGTGACCAGGAACTGATCCGTCAGGGTACCGACTACAAACACTGGCCGCAGA
TCGCTCAGTTCGCTCCGTCTGCTTCTGCTTTCTTCGGTATGTCTCGTATCGGTATGGAAGTTACCCCGTC
TGGTACCTGGCTGACCTACACCGGTGCTATCAAACTGGACGACAAAGACCCGAACTTCAAAGACCAGGTT
ATCCTGCTGAACAAACACATCGACGCTTACAAAACCTTCCCGCCGACCGAACCGAAGAAAGACAAAAAGA

AGAAAGCTGACGAAACCCAGGCTCTGCCGCAGCGTCAGAAGAAGCAGCAGACCGTTACCCTGCTGCCGGC
TGCTGACCTGGACGACTTCTCTAAACAGCTGCAGCAGTCTATGTCTTCTGCTGACTCTACCCAGGCTTAA

N_Genewiz

ATGAGCGATAACGGCCCGCAGAATCAGCGCAACGCGCCGCGCATTACCTTTGGCGGTCCGAGCGATAGCA
CCGGCAGCAACCAAAACGGAGAACGTAGCGGCGCGCGTAGCAAACAACGTCGTCCGCAAGGCCTGCCGAA
CAACACCGCGAGCTGGTTTACCGCGCTGACGCAGCATGGCAAAGAAGATCTGAAATTTCCGCGCGGCCAA
GGCGTGCCGATTAACACCAACAGCAGCCCGGATGATCAGATTGGCTATTATCGCCGCGCGACCCGTCGTA
TTCGTGGTGGTGATGGCAAAATGAAAGATCTGAGCCCGCGCTGGTATTTCTATTATCTGGGCACCGGCCC
GGAAGCGGGCCTGCCGTATGGCGCGAACAAAGATGGCATTATTTGGGTGGCGACCGAAGGCGCGCTGAAC
ACCCCGAAAGATCATATTGGTACCCGTAACCCGGCGAACAACGCGGCGATTGTGCTGCAACTGCCGCAAG
GCACCACCCTGCCGAAAGGCTTCTATGCCGAAGGTTCGCGCGGAGGGAGCCAAGCGTCATCACGCAGCAG
CAGCCGTTCGAGGAACAGCAGCCGCAACAGCACTCCTGGTAGTTCACGTGGCACATCACCGGCACGTATG
GCAGGCAATGGCGGTGACGCAGCGCTGGCGCTGCTGCTACTGGATCGCCTGAATCAGCTGGAAAGCAAAA
TGAGCGGCAAAGGTCAGCAGCAGCAAGGCCAAACCGTAACGAAGAAAAGCGCGGCGGAAGCGAGCAAAAA
ACCGCGTCAGAAACGCACCGCGACCAAAGCGTATAACGTGACCCAAGCGTTTGGCCGCCGCGGCCCGGAA
CAGACCCAAGGCAACTTTGGCGATCAAGAACTGATTCGCCAAGGCACCGATTATAAACATTGGCCGCAGA
TTGCGCAGTTTGCGCCGAGCGCGAGCGCGTTGTTTGGCATGAGCCGCATTGGCATGGAAGTGACCCCGAG
CGGCACCTGGCTGACCTATACCGGCGCGATTAAACTGGATGATAAAGATCCGAACTTTAAAGATCAAGTG
ATTCTGCTGAACAAACATATTGATGCGTATAAAACCTTTCCGCCGACCGAACCGAAGAAAGATAAAAAGA
AGAAAGCGGATGAAACCCAAGCGCTGCCGCAGCGTCAGAAGAAACAGCAGACCGTTACACTGCTGCCGGC
GGCGGATCTGGATGATTTTAGCAAACAGCTGCAGCAGAGCATGAGCAGCGCGGATAGCACCCAAGCGTAA

N_CH

ATGAGTGACAACGGGCCGCAAAACCAAAGAAACGCACCTAGAATAACGTTCGGGGGGCCGTCTGACAGTA
CGGGGTCTAACCAGAACGGGGAGAGGTCTGGGGCTAGATCTAAGCAAAGACGGCCGCAAGGGCTACCGAA
CAACACGGCTTCCTGGTTCACTGCCCTAACACAACACGGGAAGGAGGACCTAAAGTTCCCCAGAGGGCAA
GGGGTCCCCATAAACACTAACTCCTCCCCAGACGACCAAATAGGGTACTACAGGCGGGCTACAAGAAGAA
TAAGAGGGGGGGACGGGAAGATGAAGGACTTGTCACCACGGTGGTACTTCTACTACCTAGGAACAGGGCC
CGAGGCTGGGCTACCGTACGGGGCTAACAAGGACGGGATAATATGGGTTGCCACAGAGGGGGCCCTAAAC
ACACCCAAGGACCACATAGGGACTAGGAACCCAGCCAACAACGCCGCCATAGTACTACAACTACCACAAG
GGACAACACTACCCAAGGGATTCTACGCAGAGGGGTCAAGGGGAGGGTCTCAAGCTAGCTCAAGATCCAG
TTCTAGGTCTAGAAACTCTTCGCGGAACTCTACACCCGGGTCAAGTCGGGGGACAAGCCCCGCCCGGATG
GCAGGGAACGGGGGGGACGCTGCTCTAGCCCTATTGCTGCTAGACAGGCTAAACCAGCTAGAGTCTAAGA
TGTCAGGGAAGGGGCAACAACAACAAGGGCAAACAGTAACAAAGAAGTCTGCTGCAGAGGCCTCCAAGAA
GCCACGGCAAAAGAGGACAGCTACGAAGGCCTACAACGTCACGCAAGCATTCGGGCGGAGGGGGCCCGAG
CAAACTCAAGGGAACTTCGGGGACCAGGAGCTAATAAGACAAGGGACGGACTACAAGCATTGGCCCCAAA
TAGCCCAATTCGCACCGTCCGCTAGTGCTTTCTTCGGGATGTCTAGAATAGGGATGGAGGTCACACCATC
TGGGACTTGGCTAACATACACAGGGGCTATAAAGCTAGACGACAAGGACCCCAACTTCAAGGACCAAGTA
ATACTATTGAACAAGCACATAGACGCATACAAGACATTCCCACCAACGGAGCCAAAGAAGGACAAGAAGA
AGAAGGCCGACGAGACGCAAGCACTACCTCAGAGGCAGAAGAAGCAGCAAACGGTAACGCTACTACCAGC
CGCCGACCTAGACGACTTCTCCAAGCAACTACAACAAAGCATGTCTTCCGCAGACTCTACACAGGCATAA
